# A dissociation between the use of implicit and explicit priors in perceptual inference

**DOI:** 10.1101/2023.08.18.553834

**Authors:** Caroline Bévalot, Florent Meyniel

## Abstract

The brain constantly uses prior knowledge of the statistics of its environment to shape perception. These statistics are often implicit (not directly observable) and gradually learned from observation; but they can also be explicitly communicated to the observer, especially in humans. In value-based decision-making, these priors are treated differently depending on their implicit or explicit origin creating the “experience-description gap”. Here, we show that the same distinction also applies to perception. We created a pair of categorization tasks with implicit and explicit priors respectively, and manipulated the strength of priors and sensory likelihood within the same human subjects. Perceptual decisions were influenced by priors in both tasks, and subjects updated their priors in the implicit task as the true statistics changed. Using Bayesian models of learning and perception, we found that the weight of the sensory likelihood in perceptual decisions was highly correlated across subjects between tasks, and slightly stronger in the implicit task. By contrast, the weight of priors was much less correlated across tasks, and it increased markedly from the explicit task to the implicit task. The same conclusion holds when using the subjects’ reported priors. Model comparison also showed that different computations underpinned perceptual decisions depending on the origin of the priors. Taken together, those results support a dissociation in perceptual inference between the use of implicit and explicit priors. This conclusion could resolve conflicting results generated by the indiscriminate use of implicit and explicit priors when studying perception in healthy subjects and patients.

**Significance Statement:** The use of prior knowledge is ubiquitous in brain processes. However, this use is not always appropriate, and the weights assigned to priors in perceptual decisions are quite heterogeneous across studies. Here, we tested whether the origin of priors can partially explain this heterogeneity. Priors that are explicitly communicated (e.g., a road sign indicating the likely presence of wildlife) appear to be used differently from priors that remain implicit and are learned from experience (e.g., previous encounters with wildlife). We show that the use of implicit and explicit priors is unrelated across subjects. This dissociation may explain why previous findings on the use of explicit and implicit priors (e.g. abnormally strong or weak priors) are often inconsistent.

## Introduction

The use of prior knowledge reflecting the statistics of the environment is ubiquitous in cognition. In perception, studies over more than a century (Helmholtz H., 1867; Ramachandran, 1988) have shown that the brain uses prior knowledge to compensate for the frequent poverty and ambiguity of the data provided by the sensory organs (called sensory likelihood). These priors shape perceptions (Summerfield & de Lange, 2014), but also decisions based on the prospect of some reward (Rescorla & Wagner, 1972; Sutton & Barto, 2018), expectations of future events (Meyniel et al., 2016; Téglás et al., 2011), language acquisition and processing (Griffiths & Tenenbaum, 2006; Heilbron et al., 2022), and more abstract forms of reasoning (Gigerenzer, 2008; Tenenbaum et al., 2011).

Humans can make a more or less appropriate use of these priors when combined with the sensory likelihood. This combination controls a trade-off between information that is available locally (in time and space, corresponding to the likelihood) and information provided by a global context (Knill & Richards, 1996; Wei Ji Ma, Konrad Kording, Daniel Goldreich, 2021). To illustrate, on a road they travel frequently, drivers may rely on previous encounters with wildlife (this is the global context) to identify a deer crossing when only a faint silhouette is visible ahead (the likelihood). If the prior is too strong, drivers may have the illusion that a deer is present; if the prior is too weak, they may fail to detect it in time. In the general population, the weight of priors is sometimes reported to be just right on average (Ernst & Banks, 2002; Girshick et al., 2011; Körding & Wolpert, 2006; Stocker & Simoncelli, 2006), but also sometimes too weak (Ackermann & Landy, 2015; Kahneman & Tversky, 1972) or too strong (Eldar et al., 2021; Singletary et al., 2021).

An understanding of the strikingly different weights that participants reportedly assign to priors in different studies is still lacking. One possibility is that the origin of priors may partly explain these differences (Angeletos Chrysaitis & Seriès, 2023). Here, we explored a distinction between explicit priors (which are explicitly communicated to the observer) and implicit priors (which are latent and must be learned based on experience (Behrens et al., 2007; Gershman & Niv, 2010). The former are particularly common in humans (De Lange et al., 2013; Diaconescu et al., 2020). To illustrate, tourists do not have the prior knowledge of local drivers (an implicit prior), but they can be informed of the presence of wildlife by a road sign (an explicit prior).

There are well-known examples in which the weight of priors is different depending on their implicit/explicit origin. In the reward-based decision-making literature, this difference is known as the “experience-description gap” (FitzGerald et al., 2010; Garcia et al., 2021, 2023; Hertwig & Erev, 2009) (experience/description corresponding to the implicit/explicit distinction). The comparison of different experimental fields also provides suggestive and indirect evidence that the implicit/explicit distinction may correspond to a difference in the weight of priors. In perception and sensorimotor control, priors are often learned implicitly during the course of a task (Den Ouden et al., 2010; Ernst & Banks, 2002; Körding & Wolpert, 2004; Valton et al., 2019) or during development and evolution (Girshick et al., 2011; Weiss et al., 2002); in this case the use of priors often seems appropriate. By contrast, the format of information is often purely explicit in reasoning tasks, and in this case the use of priors is often reported to be more inadequate (Gigerenzer, 2008; Jardri et al., 2017; Kahneman & Tversky, 1972).

Here, we tested within the same field, namely visual perception, whether decisions are influenced differently by priors depending on their implicit/explicit origin. Previous studies in perception have yielded conflicting results when either implicit or explicit priors were used (see review by (Angeletos Chrysaitis & Seriès, 2023)). These conflicts could be due to the implicit/explicit origin of priors, but the evidence remains inconclusive because this difference has been confounded by different tasks and participants across studies. We propose to help resolve these conflicts by characterizing the use of explicit and implicit priors within the same perceptual decision-making task and group of participants.

To anticipate our results, we found differences in perceptual decisions between explicit and implicit priors. In principle, such differences can arise from different uses of the same prior value, or from a different prior value itself. Unlike explicit priors, implicit priors are, by definition, not directly observable by participants and the experimenter. The experimenter might infer that participants weigh explicit and implicit priors differently, whereas in fact the weight is the same and the value of the participant’s implicit prior differs from the experimenter’s expectation (Angeletos Chrysaitis & Seriès, 2023). Therefore, we set out to disentangle the weight of implicit priors at the perceptual decision stage from their learning. To this end, we modeled both learning and decision as Bayesian inference (Wei Ji Ma, Konrad Kording, Daniel Goldreich, 2021), and we used this overarching framework to compare the weights of implicit and explicit priors in perceptual decisions.

## Results

### Studying perceptual inference with explicit and implicit priors

We designed a set of two tasks to compare perceptual inference in the presence of explicit and implicit priors respectively (Fig. 1A). The subject’s task was identical in both cases: categorize as house or face each noisy morphed image presented in a sequence. The likelihood of the category was estimated for each image in an independent task (see Methods), and pseudo-randomly changed across trials. On a given trial, the image category was sampled with probability 0.1, 0.3, 0.5, 0.7 or 0.9 to be a house, called prior hereafter. In the first task, priors about the category were provided explicitly before each image in the form of pictograms. Subjects were instructed that the relative size of the house and face pictograms denoted the prior value. Subjects’ reports of the prior values corresponding to each pictogram tightly correlated with their true values (Fig S1.A, mean ρ=0.76, s.e.m.=0.025, Cohen’s d=1.98, t=29.55, Wilcoxon <10^−35^), demonstrating a good understanding of the pictograms. In the second task, such explicit cues were not provided, but subjects were instructed that the latent prior was constant in uncued blocks of trials, making it possible for them to learn the prior value. To make sure that priors were strictly explicit in the first task and not implicitly informed by past images, the image categories (and priors) were randomly changed across trials. Therefore the key difference between the two tasks is the type of prior: explicit or implicit.

**Figure 1.**
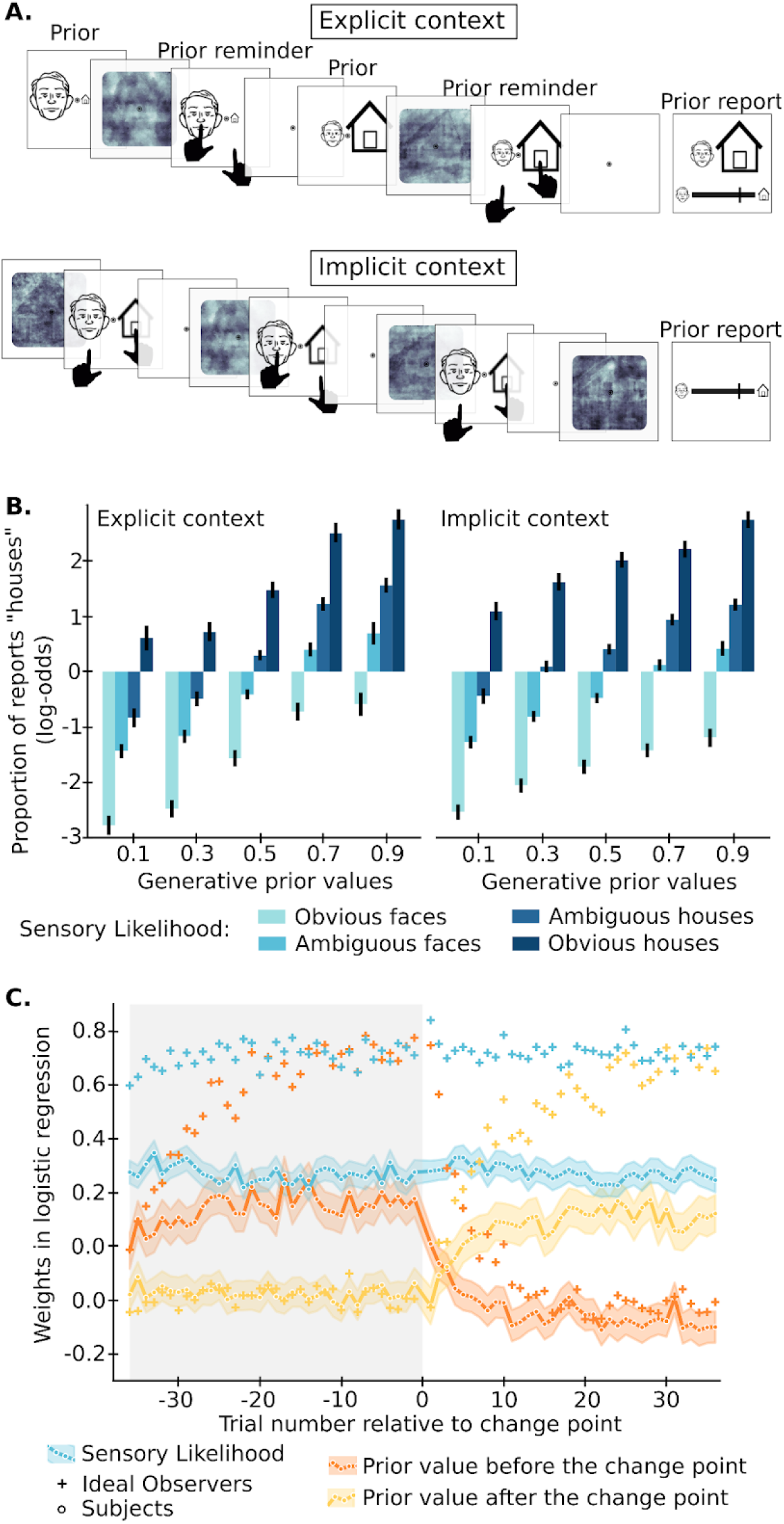
Both explicit and implicit priors influence perception. A. Task: subjects categorized each noisy image as face or house. The sensory likelihood conveyed by each image was measured in an independent group of subjects. Priors correspond to the prior probability of the latent category (face or house) on each trial. In the explicit task, priors changed from trial-to-trial and were communicated explicitly as the relative size of pictograms presented before each image (and reminded to subjects at the response stage). In the implicit task, priors were not communicated but they remained constant within uncued blocks of trials, making it possible to learn their value from the sequence of images. Subjects were occasionally asked to report the prior value. B. Average proportion of answers (plotted in log-odds) sorted by true prior values and bins of sensory likelihood. C. Logistic regression of priors and sensory likelihood onto choices, locked onto change points in the implicit task, in subjects and the BAYES-OPTIMAL model. Learning results in a change in the weights of the priors. In B-C, error bars correspond to 95% confidence intervals.

To characterize computationally perceptual inference in these two tasks, we used a Bayesian observer model. At the decision step, Bayesian inference prescribes that the posterior probability of the latent category given the image should be the product of the prior and the likelihood normalized by the image probability. This relationship is advantageously simpler when expressed in log-odds: the posterior log-odds is the sum of the prior log-odds and likelihood log-odds (see Methods, eq. 3). Subject responses showed clear effects of the generative priors and the sensory likelihood in both tasks (Fig. 1B). To quantify those effects, we estimated a logistic regression of each subject’s choices using the log-odds of generative priors and of sensory likelihood as predictors (as in Bayesian inference). The prior and sensory likelihood both influenced subjects’ choices significantly (in the explicit task, prior: mean weight= 0.82, s.e.m.=0.06, Cohen’s d=0.81, t=13.54, Wilcoxon <10^−42^; likelihood: mean weight=0.53, s.e.m.=0.02, Cohen’s d=1.26, t=21.03, Wilcoxon<10^−40^, in the implicit task, prior: mean weight=0.36, s.e.m=0.02, Cohen’s d=1.34, t =22.33, Wilcoxon <10^−41^; likelihood: mean weight=0.64, s.e.m.=0.03, Cohen’s d=1.38, t=22.94, Wilcoxon<10^−42^, tests against 0).

Data analysis based on generative prior values is relevant when they are communicated explicitly, but questionable when they remain implicit. To illustrate, it is impossible to know the new generative prior value when it has just changed in the implicit task. Therefore, a model for learning the prior based on previous images in the implicit task would be valuable to compare the two tasks. Bayesian inference can again be used to this end, by inverting the generative process of the sequence of images (see Methods). Such inference (corresponding to the BAYES-OPTIMAL model hereafter) defines a benchmark for the learning of priors, and for the use of prior and likelihood in the perception decision. The BAYES-OPTIMAL model illustrates that generative prior values can only provide a limited account of behavior in the implicit task. A trial-by-trial logistic regression analysis locked on change points in prior values (with the log-odds of prior and likelihood as predictors) showed that simulated choices of the BAYES-OPTIMAL model are not influenced by the new generative prior value immediately after a change point. Instead the effect of the new generative prior gradually builds up as the learned prior is being updated. Those dynamics are a signature of prior learning, and they are clearly visible in subjects too (Fig 1C).

### Subject-level modeling of perceptual decisions and learning

Human participants may depart from the BAYES-OPTIMAL model in different ways. At the perceptual decision stage, they may over or underweight the prior and the sensory likelihood or have responses biases. Fig. 1B already reveals such differences. At the learning stage, subjects may assume that the generative priors change more or less often than they actually do (Nassar et al., 2012), or have an exacerbated, dampened or biased use of the sensory likelihood. We designed a parameterized version of the Bayesian observer (the BAYES-FIT-ALL model) to characterize those departures from the BAYES-OPTIMAL model. We verified the recoverability of parameter estimates (see Fig S2A) and fitted the model to the choices of each subject separately in the two tasks (see Methods; note that in the explicit task, both models use the generative priors that are communicated explicitly). We used the mean cross-validated log likelihood of choices (cvLL) as a performance metric to compare models with different degrees of freedom. The (parameter-free) BAYES-OPTIMAL model captured subjects’ behavior better than chance (cvLLchance=log(0.5)=-0.69) in the task with explicit priors (cvLL=-0.51, s.e.m=0.01, Cohen’s d=1.08, t=16.11, Wilcoxon<10^−27^, test of the difference from chance) and in the task with implicit priors (cvLL=-0.50, s.e.m=0.01, Cohen’s d=1.31, t=19.56, Wilcoxon<10^−31^, test of the difference from chance). As expected based on previous studies (Wilson & Collins, 2019), by taking into account subject-specific aspects of perceptual inference the BAYES-FIT-ALL model better accounted for the subjects’ choices in the explicit task (cvLL=-0.44, s.e.m.=0.008; Cohen’s d=0.96, t=10.31, Wilcoxon<10^−23^, test of the difference with BAYES-OPTIMAL) and the implicit task (cvLL=-0.44, s.e.m.=0.007, Cohen’s d=0.85, t=12,72, Wilcoxon<10^−31^).

Some deviations from the BAYES-OPTIMAL model may be more critical than others to account for the subject’s choices. We tested simpler versions of the BAYES-FIT-ALL model, fixing some learning parameters to their optimal values and fitting the perceptual decision parameters. The model with optimal learning parameters (thereafter BAYES-FIT-DECISION) performed slightly better than models that also included free parameters at the learning stage (Fig S2B), indicating that deviations in the perceptual decision parameters are more critical than deviations in the learning parameters. To be conservative, in the main text below we report results based on the BAYES-FIT-ALL model that estimates each subject’s perceptual decision parameters while taking into account their specific learning parameters, and we report similar conclusions with alternative analyses in Supplementary Fig. 3.

### A dissociation between the use of explicit and implicit priors

We compared choices in the two tasks using a logistic regression with the (log-odds of the) posterior probability about the image category computed by the BAYES-FIT-ALL model (thereby taking into account subject-specific parameters in both perceptual decision and learning). Subjects’ choices followed this posterior more when priors were implicit than when they were explicit (Fig 2A, mean inter-task difference=-0.23, s.e.m.=0.019, Cohen’s d=-0.84, tvalue=-12.5, Wilcoxon p<10^−22^). This difference could be driven by choices being more influenced by the sensory likelihood or priors (or both) in the case of implicit priors. A logistic regression modeling the (log-odds of the) likelihood and the prior (separately from posteriors) revealed that subjects’s choices were more influenced by both the prior (mean inter-task difference=-0.42, s.e.m.=0.084, Cohen’s d=-0.33, t=-5.00, Wilcoxon p <10^−9^) and the likelihood (mean inter-task difference=-0.22, s.e.m.=0.019, Cohen’s d=-0.80, t=-11.92, Wilcoxon p<10^−23^) when priors are implicit. The difference was slightly more pronounced for the prior than the likelihood (mean difference of difference=0.20, s.e.m.=0.092, Cohen’s d=0.14, t=2.16, Wilcoxon p=0.003). Interestingly, the standard error of the mean of the paired difference was 4 times larger for the prior than the likelihood, indicating that the weight of the likelihood was more conserved across tasks than the weight of the prior. To further characterize this dissociation, we compared the between-subject correlations of weights between tasks and found a strong correlation for the likelihood (spearman rho=0.72, slope=0.75, t=15.53, p<10^−34^; Fig 2E) and only a weak correlation for the prior (spearman rho=0.145, slope=0.089, t=2.18, p=0.030; Fig 2D; significance of the difference: p<0.0001 bootstrap).

**Figure 2.**
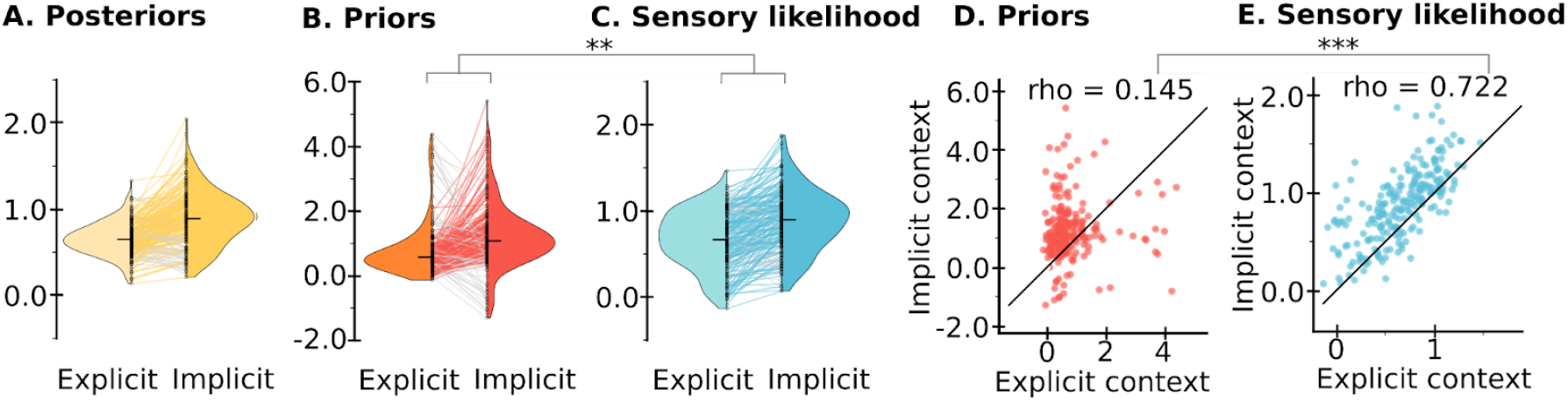
Dissociation between contexts in the use of priors, rather likelihood. Between-task comparison of the weights of the BAYES-FIT-ALL posterior probability of the category (A), the prior (B), and the sensory likelihood (C) onto subjects’ choices (one line is one subject). Weights were estimated in a logistic regression with the log-odds predictor values, so that the optimal weight is 1 in all cases (see eq. 3 in Methods). Weights in B and C were estimated in the same regression model; note that the task-difference is larger for the prior. D-E. Small correlation of prior weights (D) and strong correlation of sensory likelihood weights (E) between tasks (one dot is one subject, the black line is the identity) ** : p<0.005 ; ***: p<0.0005

### Robustness of the dissociation

The fact that the weights of the prior are much less correlated between tasks than the weights of the likelihood was robust across models: the same dissociation was found when using the generative priors, and the optimally-estimated priors (BAYES-OPTIMAL), instead of the priors based on subject-specific learning parameters (BAYES-FIT-ALL), see Fig S3.

However, a concern remains: the dissociation could be due to the inability of these models to capture the priors used by subjects in the implicit task. To rule out this possibility, we asked subjects, occasionally during the task, to report the value of the current prior on a scale (see Methods). Their reports corresponded remarkably well to the BAYES-FIT-ALL priors at the subject-level (the correlation was significant at p<0.05 in 90% of subjects) and the group-level (Fig 3.A; mean ρ=0.53, s.e.m.=0.026, Cohen’s d=1.38, t=20.67, Wilcoxon p<10^−31^). In fact, the BAYES-FIT-ALL prior proved superior to the reported prior to predict the subjects’ choice on the first trial after the report (Figure 3.B, mean difference=-0.15, s.e.m.=0.03, Cohen’s d=-0.33, t=-4.95, Wilcoxon p<10^−8^). The weight of the reported prior in the case of implicit task was also largely uncorrelated to the weight of the prior in the case of explicit task (Figure 3.C, spearman correlation rho=0.069, slope=0.039, t=1.035, p=0.30) replicating the result obtained with model-based priors.

**Figure 3.**
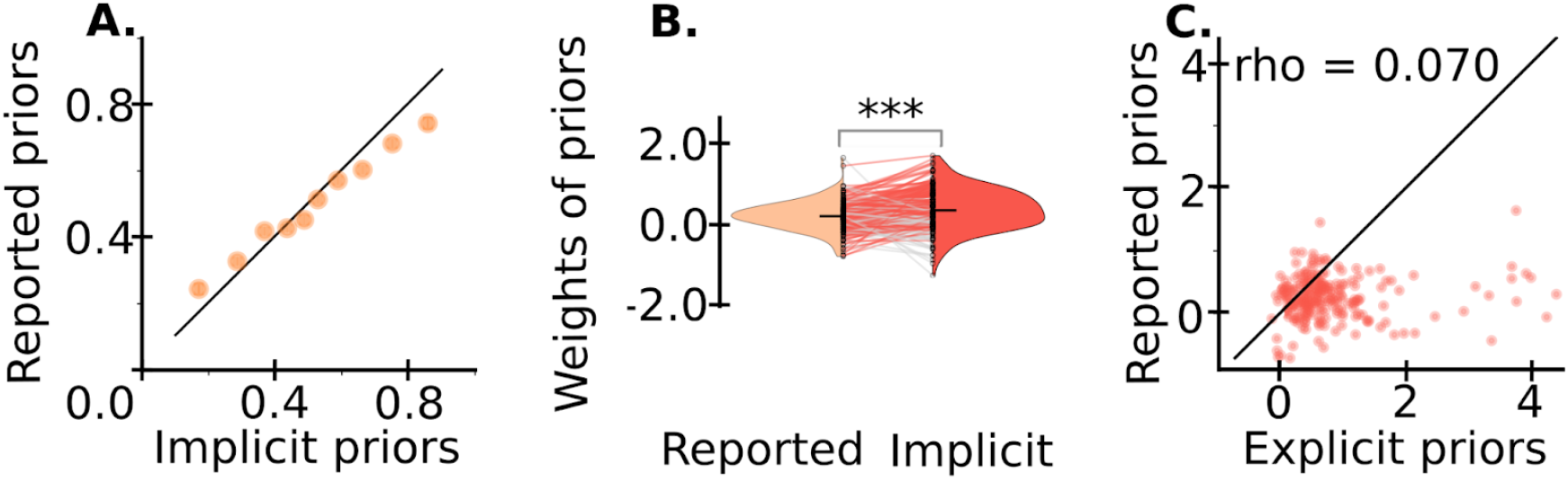
Similar dissociation with a model-free estimation of priors. A. Prior values reported by subjects correlated tightly with the implicit prior values estimated by the BAYES-FIT-ALL model (s.e.m. shown). B. Logistic regression weights of priors on the choices that immediately followed each report (one line is one subject). Note that the BAYES-FIT-ALL prior better explains the subject’s choices. C. Absence of correlation (p>0.3) between the weights of reported priors in the implicit task and the weights of priors in the explicit task (one dot is one participant). ***: p<0.0005

### Different computations with explicit and implicit priors

So far, we have characterized, within the same model, a dissociation in the weights of priors depending on their type. We now explore another potential facet of this dissociation: the use of different computations with explicit and implicit priors. The lower weights of the posterior probability of the image category computed with the Bayesian model and the lower weights of the components of this posterior (prior and likelihood) when priors are explicit could suggest a reduced adherence to Bayesian principles in this case. To test this hypothesis, we considered a series of heuristic models: a model that combined linearly the prior and likelihood (instead of the non-linear Bayesian combination, which is linear only in log-odds) or adds an interaction term between prior and likelihood in an attempt to mimic the Bayesian combination while remaining different (we verified model recovery, see Fig S4). We also considered simpler heuristic models with only the prior or the likelihood. We compared these heuristic models to the BAYES-FIT-DECISION model: all models use the BAYES-OPTIMAL prior and differ only in the use of this prior and the likelihood.

In the case of implicit priors, the best model was the BAYES-FIT-DECISION, the second best model was the heuristic model with the likelihood and prior without interaction – hereafter HEURISTIC model (Fig 4A, difference in cvLL: 0.00045, s.e.m.=0.00045, Cohen’s d=0.067, t=1.007, Wilcoxon p=0.22). In the case of explicit priors, the best model was the HEURISTIC one, and the BAYES-FIT-DECISION one was only the third-best model (difference in cvLL:-0.0024, s.e.m.=0.00013, Cohen’s d=-0.12, t=-1.82, Wilcoxon p=0.00047). The difference in cvLL between these two models was significant between tasks (difference of difference of cvLL: 0.0028, s.e.m.=0.0013; Cohen’s d=0.14; t=2.12, Wilcoxon p<10^−4^), indicating that different computations underlie perceptual inference in the two tasks.

**Figure 4.**
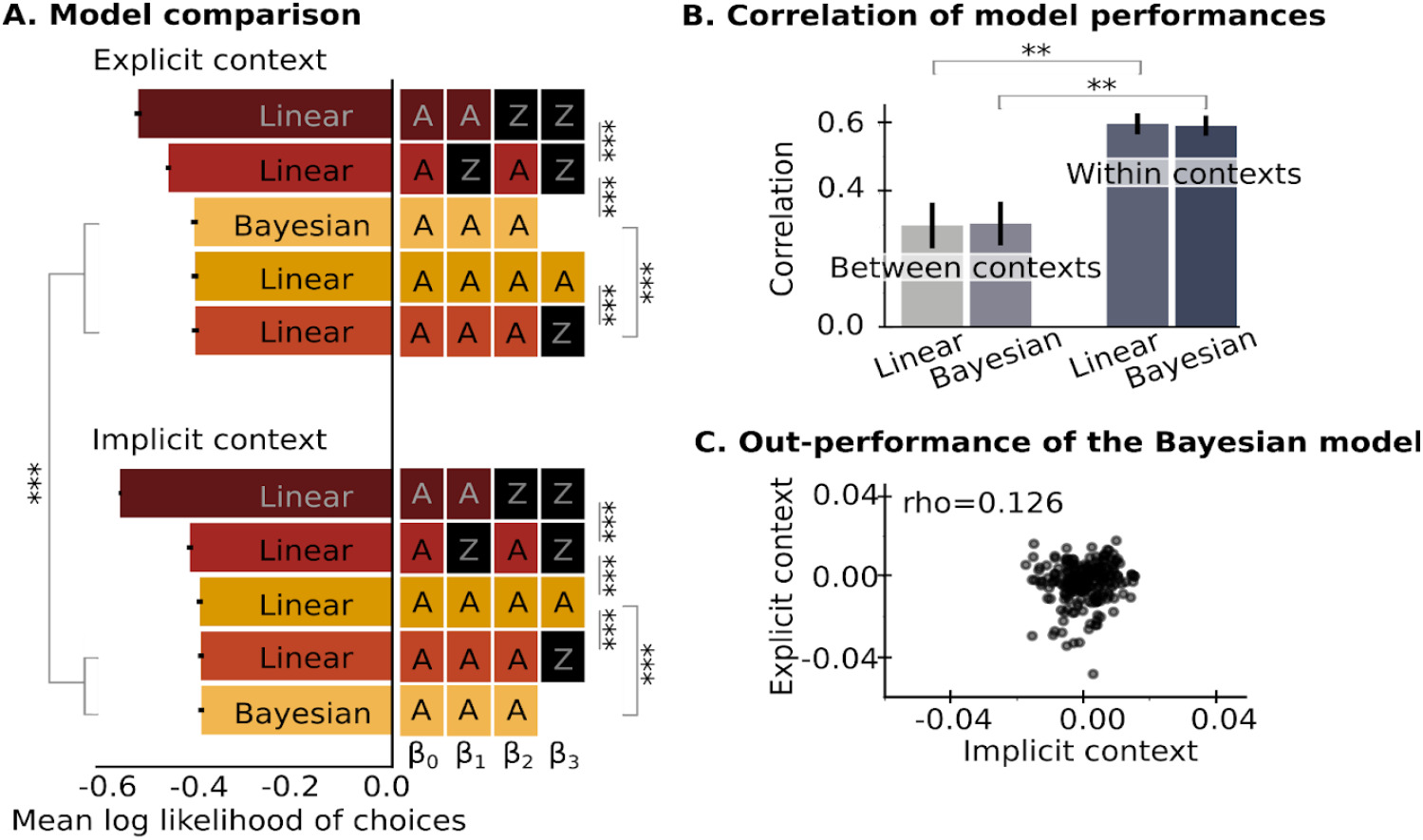
Different computations in perceptual inference depending on the prior type. A. Median performance (defined as mean cross-validated log likelihood of choices) across subjects for each model (chance level corresponds to log(0.5)=-0.69), error bars correspond to 95% CI. The best model corresponds to the model with highest log likelihood. Black cells correspond to a parameter fixed to 0, the other parameters are adjusted. Note the interaction between model types (linear vs. Bayesian) and tasks within the best models. B. Correlation of performance was estimated within tasks (using folds) and between tasks for each model (Bayesian, and linear without interaction). Bars and error bars indicate the mean and 95 CI of the bootstrapped estimation. C Correlation of the difference in model performance across tasks (Bayesian minus linear; one dot is one participant). **: p<0.005, ***: p<0.0005

In order to further test whether computations are different depending on the prior type, we estimated the between-subject correlation in model performance, within and between tasks (see Methods). If computations are subject-specific and independent of the prior type, we expect similar between- and within-task correlations; by contrast, if computations depend on the prior type, we expect a larger correlation within tasks than between tasks. Within-task and between-task correlations were markedly different, for the BAYES-FIT-DECISION model (between-task correlation ρ=0.32, slope=0.20, t=4.99, p<10^−6^, 95% confidence interval of the bootstrapped mean correlation=[0.18, 0.46] and within-task correlation (mean on the folds) ρ=0.59, slope=0.47, t=11.29, p<10^−13^ in each pair of folds, 95% confidence interval of the bootstrapped mean correlation=[0.53, 0.64] and 95% confidence interval of the bootstrapped difference in mean correlation between or within-task=[0.12, 0.41]). The same was true for the HEURISTIC model (between-task ρ=0.30, slope=0.18, t=4.76, p<10^−6^, bootstrapped mean correlation 95% CI=[0.18, 0.46] and within-task (mean on the folds) ρ=0.60, slope=0.49, t=11.41, p<10^−13^ in each pair of folds, bootstrapped mean correlation 95% CI=[0.53, 0.65]; bootstrapped difference 95% CI=[0.13 ; 0.43]), see Fig 4B. Note that smaller between-task correlations could be partly driven by some subjects being better in one task than the other (which would be reflected for each model by a corresponding difference in cvLL between tasks and not, or less so, within-tasks). We thus tested whether the paired difference in cvLL between models (which reflects the preference for a type of computation) was correlated between tasks, and we found only a small correlation (spearman ρ=0.126, p=0.060, Fig 4C), which is again indicative of different computations between the two tasks.

## Discussion

We used the framework of Bayesian inference to compare the use of implicit and explicit priors in perceptual decision making. We found three main differences. First, the weights of explicit priors were smaller (and further from the optimal weight) than those of implicit priors. Second, the weights of priors were hardly correlated across tasks and individuals. In contrast, likelihood weights were more similar on average and highly correlated across tasks. Third, perceptual decisions were supported by different computations: the prior-likelihood combination was closer to Bayesian integration when priors were implicit, and closer to a simpler heuristic when they were explicit.

Our finding that perceptual decisions are more optimal when priors are implicit (rather than explicit) is consistent with previous findings. Studies in which participants are presented with explicit information have often emphasized that their behavior deviates from optimality (e.g. probability judgments or value-based decisions (Eldar et al., 2021; Hotaling et al., 2015; Kahneman & Tversky, 1972; Sides et al., 2002). In contrast, behavior has been reported to be close to the optimum when information remains more implicit and dominated by low-level processing (e.g., in perception (Girshick et al., 2011; Knill & Richards, 1996; Stocker & Simoncelli, 2006; Weiss et al., 2002) and sensorimotor control (Körding & Wolpert, 2004; Wolpert et al., 1995). However, comparisons with the optimum in these studies remain mostly qualitative, and their conclusions may reflect the perspective of the authors. Quantifying this optimality would be useful to compare different studies using the same metric. Unfortunately, direct and quantitative comparisons of decisions based on implicit and explicit priors are rare (Leptourgos et al., 2020; Thakur et al., 2021).

Our estimate of prior and likelihood weights has the advantage of being quantitative. It is also absolute rather than simply relative. Absolute quantification is possible here because our task involves categorical choices (see eq. 3). It is not possible in tasks involving continuous percepts and Gaussian distributions (a common experimental choice). The mean percept resulting from a Gaussian prior and a Gaussian likelihood is in principle the average of their means weighted by their relative precisions. Since the weighting is relative in this case, the effects of a weaker (i.e., more imprecise) prior and a stronger (i.e., more precise) likelihood are indistinguishable (Brock, 2012). Note that when categorical percepts are used (which allows estimation of absolute weights), sometimes only the relative prior-likelihood weight is estimated due to specific modeling choices (Powers et al., 2017; Schmack et al., 2017).

Differences between tasks can be quantified within the same model, but computations may in fact differ profoundly between tasks. The difference in computations that we found for perceptual decisions based on implicit and explicit priors is reminiscent of a similar difference reported for value-based decisions (Hertwig & Erev, 2009). Specifically, we found that the prior-likelihood integration was more Bayesian with implicit priors. This conclusion is based on the contrast between the Bayesian computation and a linear approximation. We emphasize that this comparison is relative; quantifying the extent to which a computation is Bayesian would require more model comparisons (Adler & Ma, 2018; Laquitaine & Gardner, 2018). For example, models of circular prior-likelihood integration have been proposed by others (Jardri et al., 2017; Leptourgos et al., 2020).

Our characterization of the differences between the use of explicit and implicit priors is at the computational level. It raises interesting possibilities in terms of their neurobiological implementation. For example, the difference in their use could arise from the fact that the neural representations of explicit and implicit priors are different. In value-based decision-making, the neural correlates of explicit and implicit prior values were measured with functional MRI and found to be indeed anatomically distinct (FitzGerald et al., 2010).

Another, non-mutually exclusive possibility is that the prior-likelihood integration recruits different brain circuits depending on the format of the priors. When priors are implicit, this integration could be handled by the interaction of statistical learning, and perceptual decision making systems (Fiser & Lengyel, 2022; Sherman et al., 2020). Such integration may operate beyond the scope of consciousness, as often reported in statistical learning (Atas et al., 2014). In contrast, when priors are explicit, a symbolic or semantic understanding is required. Therefore, the integration of explicit priors with likelihood may recruit sensory areas, associative areas such as the parietal cortex, and hubs such as the temporal cortex or the prefrontal cortex associated with semantic knowledge or conscious access (Mashour et al., 2020; Ralph et al., 2017). Thus, prior-likelihood integration may rely on different levels of processing when priors are implicit or explicit, recruiting lower or higher levels of the cortical hierarchy, respectively. Such a difference in processing levels could explain the more Bayesian integration observed with implicit priors, as lower-level processes have been hypothesized to be more Bayesian (Hansmann-Roth et al., 2021; Vul & Pashler, 2008; Yeon & Rahnev, 2020).

Our results contribute to the understanding of the basic principles of perceptual decision making in the general population. They are also relevant to the study of disorders. Perception has been extensively studied in psychiatry because the representation of priors and likelihoods, and their integration, have been hypothesized to be altered, particularly in autism (Pellicano & Burr, 2012) and schizophrenia (Valton et al., 2017). Empirical evidence remains mixed. In the case of autism, it has been suggested that the nature of prior (implicit/explicit) should be taken into account, as alterations appear to be more pronounced for implicit than for explicit priors (Angeletos Chrysaitis & Seriès, 2023). In the case of schizophrenia, experiments have used priors that are either explicit (Baker et al., 2019; Jardri et al., 2017; Speechley et al., 2010) or implicit (Salvador et al., 2022; Sanders et al., 2013; Valton et al., 2019). Evidence for altered prior-likelihood combination is also mixed in schizophrenia. For example, the weight of priors in the beads task has been reported to be abnormally strong in patients when priors are explicit (Speechley et al., 2010) and abnormally weak when they are implicit (Salvador et al., 2020). Our results in the general population showed that the weights of explicit and implicit priors are essentially unrelated, so it is expected that the alteration of the prior weights observed in one group of patients may not generalize across prior types.

We now turn to some of the limitations and open questions of our study. One weakness is that the order of the tasks was fixed across subjects, always starting with implicit priors. However, the order of the task seems unlikely to have influenced the main results. The larger prior weight in the second task could in principle be due to a gradual improvement with task exposure, but the evolution of the effects during the task actually rules out this possibility (Fig S5). Furthermore, the task order cannot explain the negligible correlation of prior weights between tasks. A second limitation is that a better communication of explicit priors could have led to larger prior weights in perceptual decisions. To mitigate this concern, we used ratings to verify that subjects correctly understood the pictograms we used to communicate priors. However, this may not be sufficient, as the format of probabilistic information is known to influence the success of its use, particularly whether it adheres to Bayesian principles in reasoning tasks (Gigerenzer & Hoffrage, 1995). Third, our results leave open the stage at which implicit and explicit priors shape perceptual decisions: they may bias the incoming sensory evidence (Urai et al., 2019) or the motor response (De Lange et al., 2013). This issue should be addressed in future studies.

In conclusion, our results suggest that explicitly or implicitly acquired priors are used differently in perceptual decisions. As a consequence, inferences made with explicit or implicit priors should not be grouped indiscriminately, and the relevant literature should be reviewed with this distinction in mind.

## Materials and Methods

### Participants

The study was approved by the local Ethics Committee (CER Paris Saclay, n°222) and participants gave their informed written consent before participating. 280 subjects (136 women and 143 men) aged between 19 and 52 (mean=31; SD=8) were recruited online (Prolific.co) and 277 completed the two tasks. We excluded subjects based on catch trials presenting strong and congruent sensory evidence and prior values (both with p(house)>0.85 or p(face)>0.85). Subjects whose proportion of aberrant answers on catch trials exceeded 2 standard deviations of the group-mean were excluded. The final sample size was 277-54=223 subjects.

### Tasks

The tasks were run on Gorilla™. The experiment was composed of two tasks (with explicit and then implicit priors). They contained respectively 300 trials and around 600 trials (mean = 597, SD = 15; variability arises from sampling the length of each stable period, see below). Each task was divided into three parts separated by self-paced breaks.

Stimuli were noisy, gray-scaled, morphed images of faces and houses. Noise was first added on images with the GNU image. Morphed images were created in the Fourier space, by computing a weighted average of the phase values of each pair of face and house images, and using the mean amplitudes of all images (all images therefore have the same frequency spectrum). The level of perceptual evidence in favor of the house/face category was estimated empirically for each image as the proportion of house/face responses in a categorization task performed by a group of 47 typical human observers who did not participate in the main experiment, see Supplementary Methods.

Each trial of the main experiment consisted in the categorization of an image presented for 150 ms. Subjects reported their responses on a keyboard with the keys ‘e’ and ‘p’ for face and house (counterbalanced across subjects), within a response window of 2000 ms. In the explicit task, each image was preceded by a cue consisting of a set of pictograms indicating the prior probability of the house/face category, presented for 1500 ms. In the explicit task, the latent prior value was stable in periods of 40 trials (±4) separated by unsignaled change point. In both tasks and on each trial, the latent category was sampled according to its prior probability, and an image corresponding to this category was sampled pseudo-randomly. To make the detection of a change point easier in the implicit task, three images with strong likelihood (p(house)>0.85 or <0.15 were presented among the first four images after each change point. In the task with explicit priors, subjects were prompted to report the prior value corresponding to each pictogram at the end of the task on a quasi-continuous rating scale. In the task with implicit priors, subjects were periodically (every 14 trials) asked to report the current (implicit) prior value. In supplementary information, we provide the instruction slides containing the cover story that we used to gamify the task and make it intuitive.

### Models

#### Bayesian perception model

Both tasks are to infer, on each trial *k*, whether the latent category *c*_*k*_ is a house (denoted *H)* or a face (*F*) based on the noisy image *I*_*k*_ and some prior about the category. We focus first on the explicit task, in which the prior value does not depend on previous images, but only on the cue. Formally, this inference amounts to estimating the posterior probability *p* (*c*_*k*_ = *H*|*I*_*k*_), which is done optimally with Bayes rule:

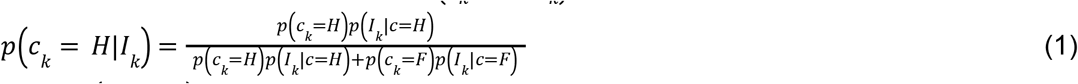

where *p* (*c*_*k*_ = *H*) is the prior probability of the latent category being a house on trial *k*, which we note θ_*k*_ for brevity below (note that*p* (*c*_*k*_ = *F*) = 1 − θ_*k*_); and*p* (*I*_*k*_ |*c* = *H*) is the sensory likelihood about the house category based on image *Ik* only. We introduce the notation 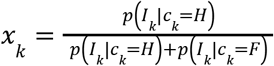 for the normalized sensory likelihood, and rewrite equation (1) more concisely as:

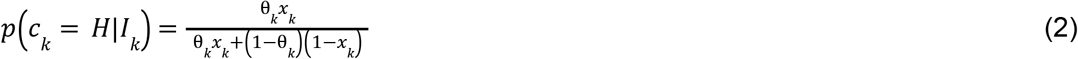

The equation is even simpler when using the log-odd transformation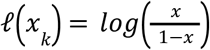:

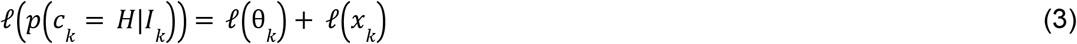

The (normalized) sensory evidence*x*_*k*_ corresponding to each image was estimated empirically in an independent group of subjects (see Supplementary Methods).

The priorθ_*k*_ was equal to its explicit value in the explicit task. To analyze the implicit task, we used in different analyses either its generative value, or its value learned based on previous images by the BAYES-OPTIMAL model or the BAYES-FIT-ALL model.

### BAYES-OPTIMAL learning model

The BAYES-OPTIMAL observer learns the priorθby updating optimally its posterior distribution after every image in the sequence (it starts with a uniform distribution before the first image). The generative process of the sequence has an interesting property (known as Markov property): if the previous prior valueθ_*k*−1_ is given, then the previous observations*I*_1_, …, *I*_*k*−1_ (denoted*I*_1:*k*−1_ below) are not informative about the current prior valueθ_*k*_. This is because here,θ_*k*_ will be equal toθ_*k*−1_ if no change point occurs (which happens with prior probability *v*, called volatility) and different otherwise; and this potential change does not depend on*I*_1:*k*−1_. This Markov property makes it possible to cast the updating process as the forward pass of a hidden markov chain (Behrens et al 2006, Meyniel, Maheu & Dehaene 2016; Gallistel et al 2014), resulting in the following iterative equation for learning the priorθ:

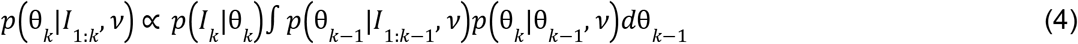

where*p* (*I*_*k*_ |θ_*k*_) is the likelihood of the current image given some prior valueθ_*k*_ ; which is obtained by marginalizing over the unknown latent category:

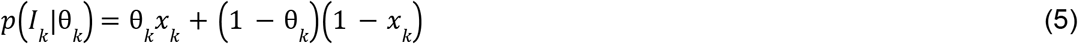

We computed the integral in eq. 4 numerically by discretization on a grid. Note that eq. 4 returns the inferred prior value after having observed*I*_*k*_. In eq. 2, the prior that is used to infer the category of image*I*_*k*_ is the prior estimated before seeing *I*_*k*_ ; it is obtained by marginalizing over the values ofθ_*k*−1_ given the previous observations*I*_1:*k*−1_ :

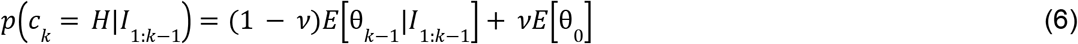

where*E*[.]denotes the expectation andθ_0_is the distribution from which new values ofθare sampled in case a change point occurs.

### BAYES-FIT-ALL model

This model differs from the BAYES-OPTIMAL with 6 free parameters that are fit to the choices of each participant. At the learning stage (eq. 4), the volatility parameter is a free parameter because the value assumed by participants may differ from the generative value. This BAYES-FIT-ALL model also allows for distortions of the likelihood function*p* (*I*_*k*_ |θ_*k*_) that can exacerbate it, dampen it, or bias it.

We considered a distortion that is affine in log-odd (Zhang & Maloney, 2012), hence yielding two additional free parameters. At the decision stage, the BAYES-FIT-ALL model has three free parameters because it considers subject-specific weightsβ_1_, β_2_ on the likelihood and prior terms in eq. 3 and an additive bias termβ_0_ (in the BAYES-OPTIMAL model,β_1_ = β_2_ = 1andβ_0_ = 0). The resulting modified eq. 3 can be fit to each participant’s choices with a logistic regression:

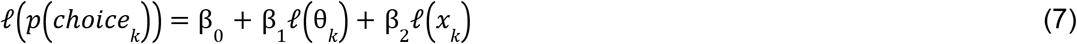

### Linear models

Those models use the prior valuesθ_*k*_ of the BAYES-OPTIMAL model and differ in the perception model: the heuristic log-odd posterior (and thus, choice probability) is simply a linear combination of the prior and sensory likelihood:

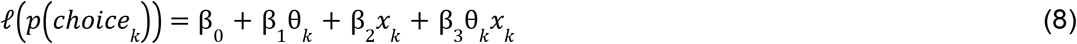

We tested different versions by fixing some parameters in {β_1_, β_2_, β_3_ to 0.

### Fitting procedure and parameter recovery

The free parameters of the BAYES-FIT-ALL model were fitted to the choices of each subject by minimizing the negative log likelihood of the subject’s choices (eq. 7). We used a Nelder-Mead algorithm implemented in Scipy’s optimize module, and we took the best fit after 50 random initializations of the parameter search to avoid local minima.

We checked the recoverability and the identifiability of parameters by applying the same fitting procedure to simulated subject choices with known parameters (Figure S2).

Depending on the task and analysis, the priors could be the explicit ones, the BAYES-OPTIMAL ones or the BAYES-FIT-ALL ones. We estimated the weights of those priors (and the sensory likelihood and bias) in each subject with logistic regression (implemented in scikit-learn with the default Ridge penalty λ = 1) in all cases for consistency. In the specific case of BAYES-FIT-ALL priors, those weights can also be estimated with the Nelder-Mead algorithm; parameter estimates were highly correlated with the logistic estimates.

### Model comparison

We compared models using their mean cross-validated log likelihood (cvLL) given each subject’s choices (Fig 4 and S1). Cross-validation penalizes models for over-fitting (Bishop & Nasrabadi, 2006), which is critical here since different models have different numbers of free parameters. We used a three-fold cross validation for each task, using as folds the sequences of images that were separated by short breaks during each task. The reported cvLL is the mean log-likelihood of choices on the left-out fold (the one not used to fit the model).

## Data and Code availability

The raw data and code to reproduce the results will be made publicly available upon publication.

## Acknowledgments

This work has been funded by the “Fondation pour la Recherche Médicale” and the Inserm. F.M. is supported by the European Research Council (ERC grant #94105).

## Supplementary Information

### Supplementary Figures

**Supplementary Figure 1.**
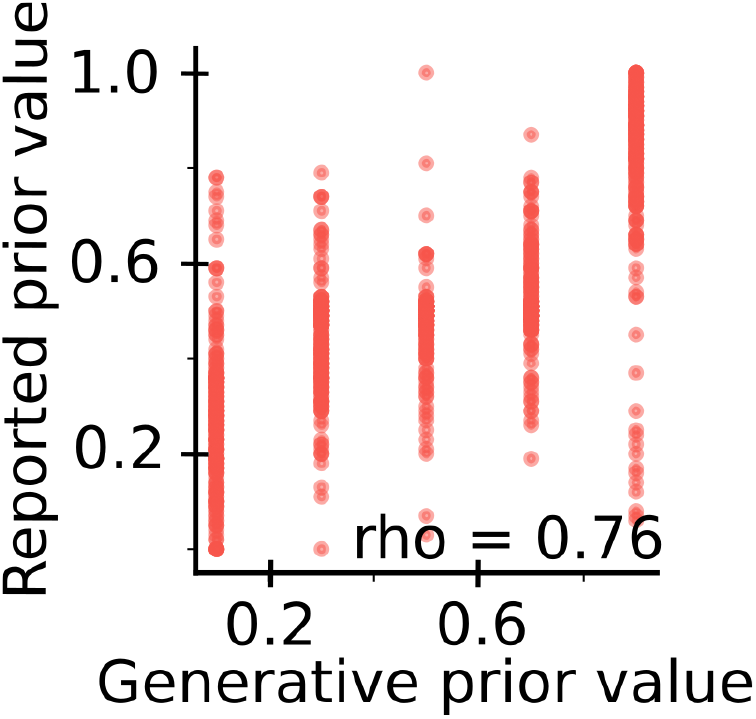
Accurate prior reports uncorrelated between tasks. Accuracy of prior reports in the explicit task (spearman coefficients of the correlation between reported prior values and generative prior values).

**Supplementary Figure 2.**
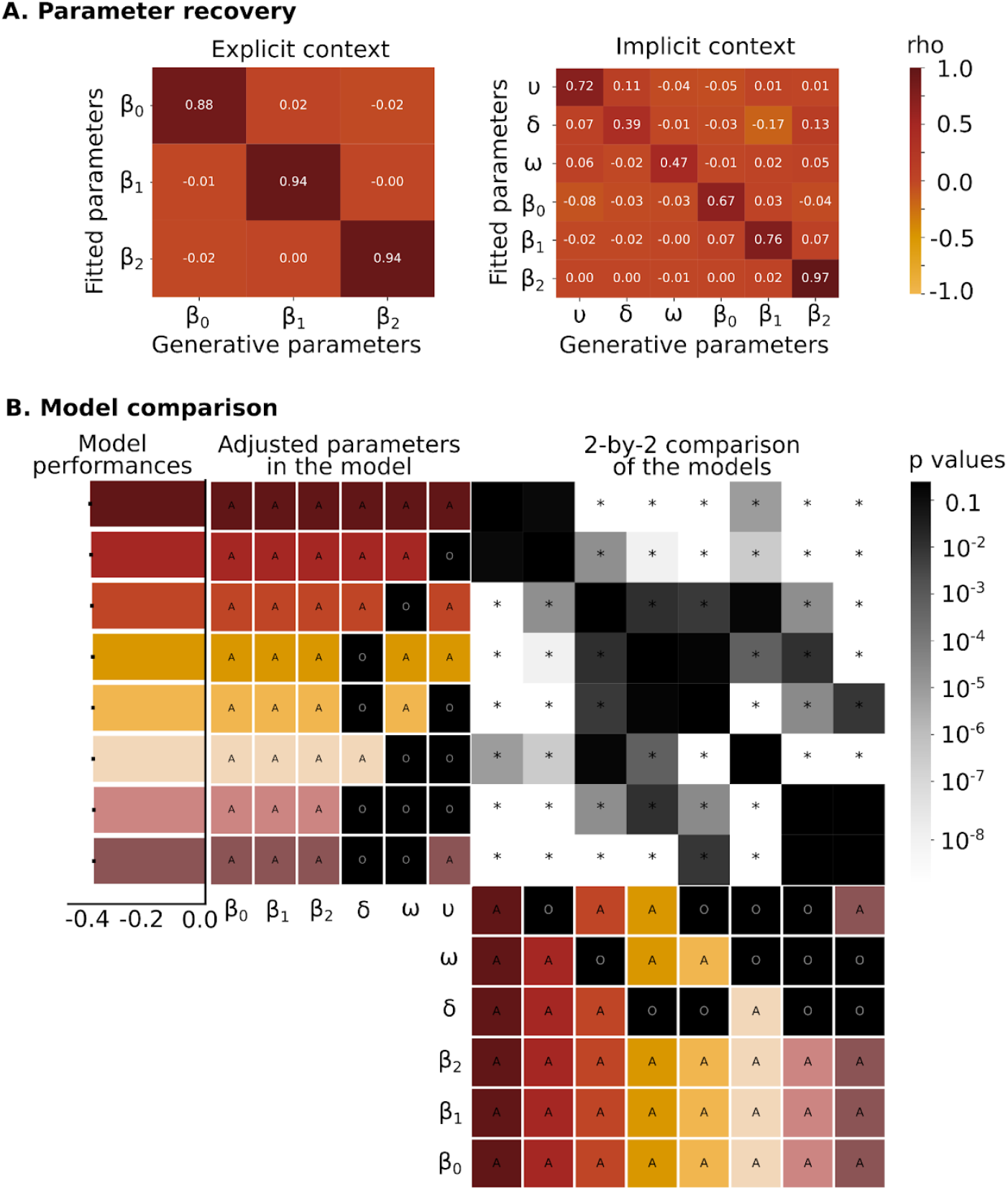
A reliable fitting procedure limiting overfitting. A. Parameter recovery of a simulated behavior. The matrix shows the spearman correlation between generative and fitted parameters in both tasks. B. Cross validated log-likelihood of Bayesian models of learning and decision with a varying number of learning free parameters. The best model corresponds to the model with highest log likelihood. U: volatility, ω: weight of sensory likelihood in the learning, δ: bias of sensory likelihood in the learning, β0: bias in sensory likelihood in the decision, β1: weight of priors in decision, β2: weight of sensory likelihood in the decision * : p <0.05

**Supplementary Figure 3.**
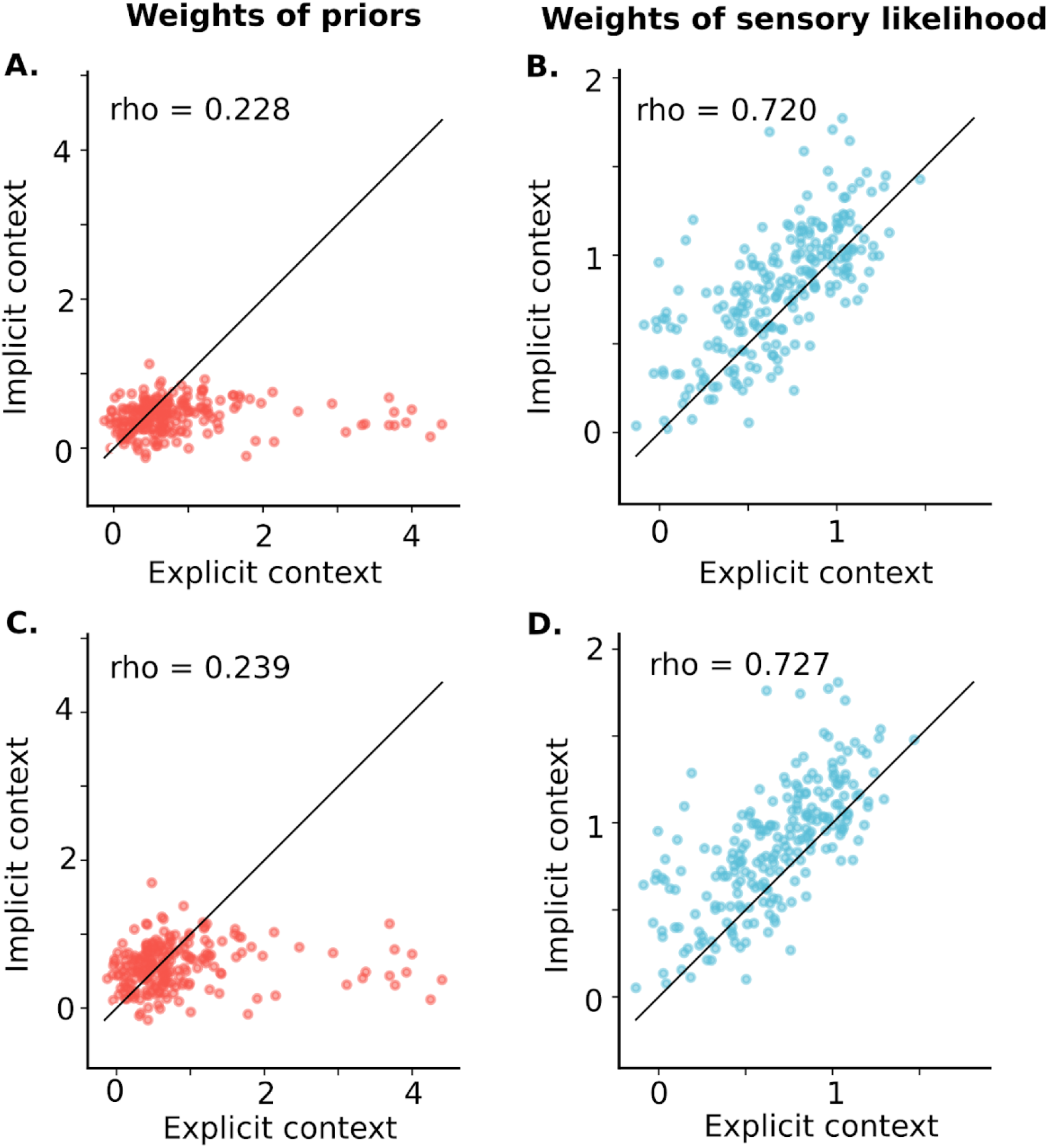
A dissociation robust to modeling choices. A. and B. Dissociation with generative priors in both tasks. C. and D. Dissociation with optimal priors in the implicit task.

**Supplementary Figure 4.**
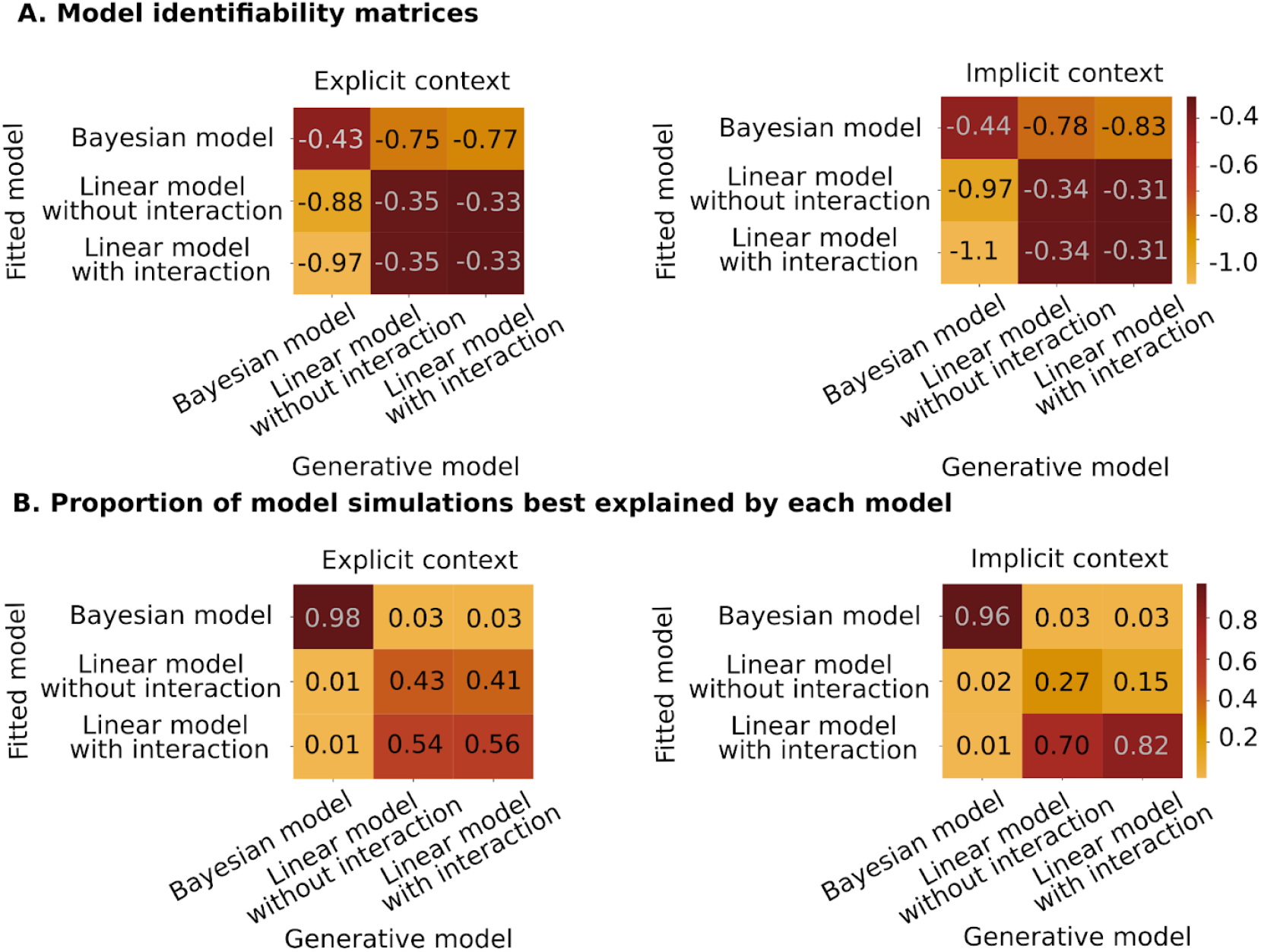
Identifiability of the Bayesian and Linear models. A. Model performances for behavior simulated with linear and Bayesian models (mean cross-validated log-likelihood). B. Proportion of model simulations (of choice behavior) that are best explained by each of the fitted models.

**Supplementary Figure 5.**
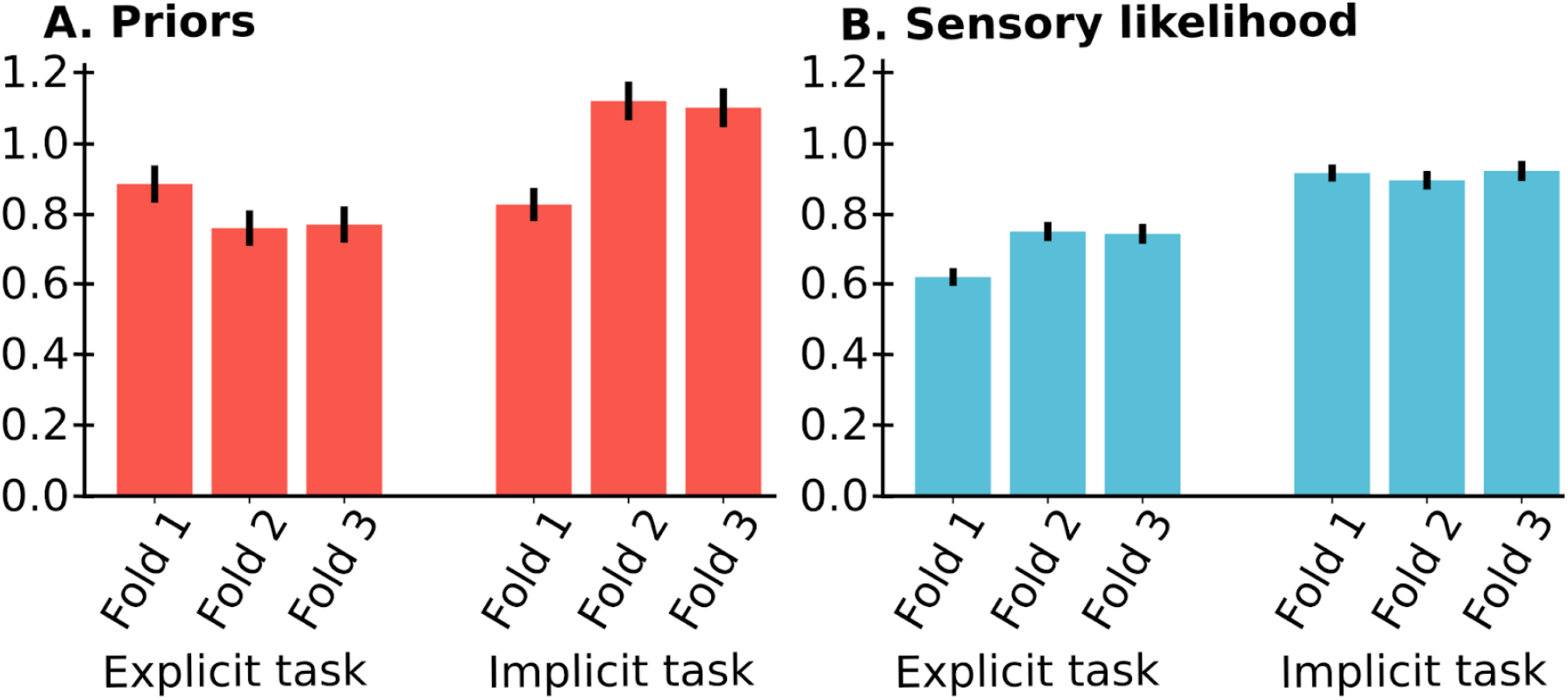
Stronger prior weights in the implicit context are not due to training. Logistic weights of prior (A.) and sensory likelihood (B) are increasing with task exposure.

#### Supplementary Methods

##### Estimation of the sensory likelihood for each image

We estimated empirically the (normalized) sensory likelihood*x*_*k*_ corresponding to each image*I*_*k*_ by asking an independent group of 47 subjects (called the rater group) to categorize each image as a house or a face. Images were presented in a random order across participants to minimize sequential effects in the average responses.

Our underlying assumption was that the group mean response for each image approximates the posterior probability of the category*p* (*c*_*k*_ = *H*|*I*_*k*_) for a typical human observer (we discuss the consequence of our approximations below). We used Bayes rule to relate*x*_*k*_ (note that it is the *normalize* sensory likelihood) to this posterior:

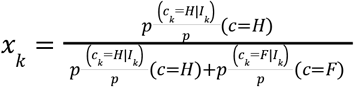

We further assumed that*p*(*c* = *H*) = *p*(*c* = *F*) = 0. 5because the sequence of images presented to raters had an equal proportion of noisy images generated dominantly from a house or a face image. In this case,*x*_*k*_ s simply equal to the average response, and constitutes our empirical estimation of the sensory likelihood.

We now discuss the consequences of our assumptions. First, If the rater group actually has unequal priors (e.g. a priori preference for faces), this will bias our estimate of*x*_*k*_. But note that such a bias would simply be additive after a log-odd transform, this bias can thus be captured as β_0_in eq. 3 (perceptual decision stage); and as the bias term for distorting the likelihood function in the learning stage (eq. 4) of the BAYES-FIT-ALL model. Second, we assumed that raters sample their categorical response from the posterior (a response strategy known as probability-matching), so that the average response approximates the posterior. If subjects have a bias in this sampling process (e.g. they report one category more than another, or they distort the posterior, e.g. with a softmax response strategy), then again those biases will be captured in the BAYES-FIT-ALL model at the decision stage (β_0_,β_2_in eq. 3) and in the parameterized distortion of the likelihood function during learning (eq. 4). Therefore, our approach for estimating the sensory likelihood remains valid when the BAYES-FIT-ALL model is used even when the above assumptions are unmet.

##### Task instructions

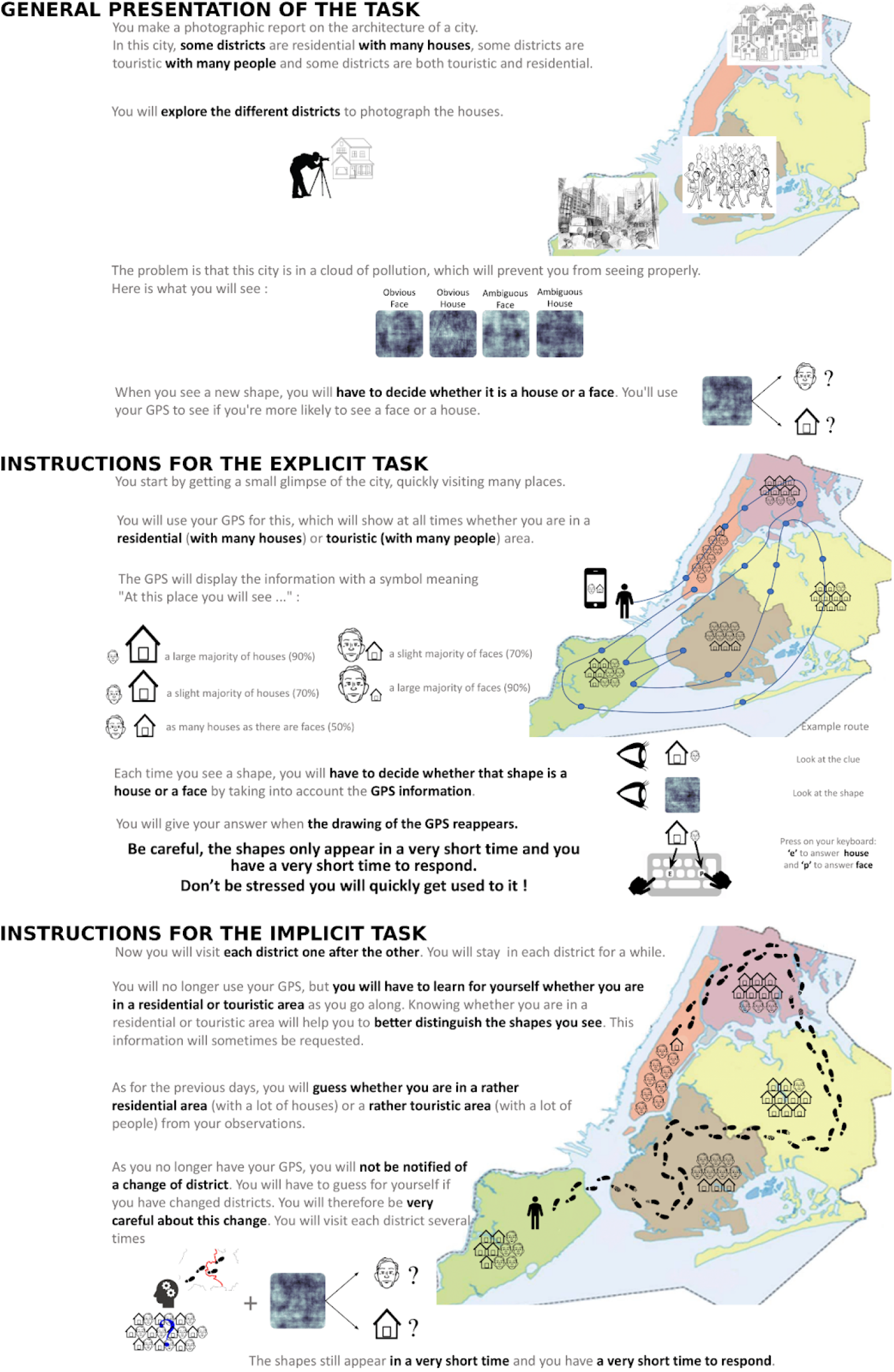

